# Functional and Pharmacological Evaluation of a Novel *SCN2A* Variant Linked to Early-onset Epilepsy

**DOI:** 10.1101/2020.04.03.024489

**Authors:** Scott K. Adney, John J. Millichap, Jean-Marc DeKeyser, Tatiana Abramova, Christopher H. Thompson, Alfred L. George

## Abstract

**Objective:** We identified a novel *de novo SCN2A* variant (M1879T) associated with infantile-onset epilepsy that responded dramatically to sodium channel blocker antiepileptic drugs. We analyzed the functional and pharmacological consequences of this variant to establish pathogenicity, and to correlate genotype with phenotype and clinical drug response.

**Methods:** The clinical and genetic features of an infant boy with epilepsy are presented. We investigated the effect of the variant using heterologously expressed recombinant human Na_V_1.2 channels. We performed whole-cell patch clamp recording to determine the functional consequences and response to carbamazepine.

**Results:** The M1879T variant caused disturbances in channel inactivation including substantially depolarized voltage-dependence of inactivation, slower time course of inactivation, and enhanced resurgent current that collectively represent a gain-of-function. Carbamazepine partially normalized the voltage-dependence of inactivation and produced use-dependent block of the variant channel at high pulsing frequencies. Carbamazepine also suppresses resurgent current conducted by M1879T channels, but this effect was explained primarily by reducing the peak transient current. Molecular modeling suggests that the M1879T variant disrupts contacts with nearby residues in the C-terminal domain of the channel.

**Interpretation:** Our study demonstrates the value of conducting functional analyses of *SCN2A* variants of unknown significance to establish pathogenicity and genotype-phenotype correlations. We also show concordance of *in vitro* pharmacology using heterologous cells with the drug response observed clinically in a case of SCN2A-associated epilepsy.

## INTRODUCTION

Variants in the *SCN2A* gene are associated with a range of childhood epilepsies differing in severity with or without accompanying neurodevelopmental delay.^1,2^ Clinical phenotypes include benign infantile seizures, developmental epileptic encephalopathy, West syndrome, and Ohtahara syndrome. *SCN2A* encodes the pore-forming subunit of the voltage-gated sodium channel Na_V_1.2. Gain-of-function variants, which result in greater Na_V_1.2 activity than normal, are associated with neonatal and infantile-onset epilepsy (onset before 3 months of age).^3^ By contrast, loss-of-function variants are associated with later-onset epilepsy with more prominent neurodevelopmental delay as well as autism spectrum disorder.^4^ More widespread genetic testing of children with epilepsy has resulted in an explosion of new *SCN2A* variants,^5^ yet the functional consequences and clear genotype-phenotype relationships have not been determined for most variants.

The majority of Na_V_1.2 channels are concentrated in the axon initial segment (AIS) of cortical excitatory neurons. During early development, Na_V_1.2 predominates throughout the AIS, acting as an action potential initiator and summing somatodendritic inputs. As neurons mature, Na_V_1.6 (encoded by *SCN8A*) replaces Na_V_1.2 in the more distal segments of the AIS. Later in development, Na_V_1.2 contributes to the back propagation of action potentials into the somatodendritic compartment of excitatory neurons.^6–8^ Gain-of-function variants in *SCN2A* are thought to cause early-onset epilepsy by promoting excitability of cortical neurons during the developmental stage when Na_V_1.2 predominates in the AIS.^9^ Clinical studies suggest that some infantile-onset epilepsies due to gain-of-function variants are responsive to treatment with antiseizure drugs (ASDs) having a principal mechanism of action to block sodium channels, albeit in a nonselective manner.^10^

Here we report a case of infantile-onset epilepsy associated with a novel *SCN2A* variant (M1879T) of uncertain significance combined with extensive functional and pharmacological studies of the variant sodium channel. Our findings reveal a novel constellation of functional abnormalities associated with infantile-onset epilepsy, and provide a demonstration that *in vitro* pharmacological properties of a variant sodium channel correlate with the clinical response to a specific drug.

## MATERIALS AND METHODS

### Study Participant

The patient was recruited to study #2015-738 approved by the Institutional review Board of Ann & Robert H. Lurie Children’s Hospital of Chicago. Following informed consent, data was collected by parental report through a REDCap database questionnaire and review of medical records.

### Mutagenesis and heterologous expression of Na_v_1.2

The open reading frame of human Na_V_1.2 was subcloned into a custom pCMV-IRES-mScarlet vector. The M1879T mutant channel was constructed using site-directed mutagenesis PCR (Forward primer: AATACAGA<u>C</u>GGAAGAGCGATTCATGGCATCAAACCC; Reverse primer: GCTCTTCC<u>G</u>TCTGTATTCGAAGGGCATCCATCTCTCC) with Q5 polymerase (New England Biolabs, Ipswitch, MA). The neonatal construct differed from the adult by inclusion of the neonatal exon 6N instead of the adult exon. All constructs were verified by Sanger sequencing of the promoter through to the polyadenylation signal, inclusive.

HEK293T cells (ATCC, Manassas, VA) were stably transfected with the β1 and β2 sodium channel subunits under puromycin selection to generate a stable cell line using previously established methods.^11^ Cells were maintained at passages 5-15 in DMEM media supplemented with 1% Pen/Strep, 10% FBS, and puromycin (3 μg/mL). The HEK293Tβ1/β2 cell line was transiently transfected using SuperFect reagent (Qiagen, Germantown, MD) with 2 μg of WT or mutant Na_V_1.2 plasmid along with 0.25 μg of a second plasmid (pEGFP-IRES-SCN2B) for use as an additional transfection marker. After 24-48 h, transfected cells were harvested with trypsin and plated on poly-D-lysine (Sigma-Aldrich, St. Louis, MO) coated coverslips prior to recording. Only fluorescent cells were selected for whole-cell patch clamp recording.

### Electrophysiology

Whole-cell patch-clamp recordings were performed at room temperature (22-23°C). Borosilicate glass pipettes were pulled with a P-1000 (Sutter Instruments, Novato, CA), with resistances of 1-2.5 MOhm. External solution contained (in mmol/L) 140 NaCl, 10 HEPES, 4 KCl, 1 MgCl_2_, 1.8 CaCl_2_, 10 dextrose, pH 7.35 adjusted with NaOH, osmolarity 310 mOSm. Internal solution contained (in mmol/L) 10 NaF, 105 CsF, 20 CsCl, 2 EGTA, 10 HEPES, 10 dextrose, pH 7.35 adjusted with CsOH, osmolarity 300 mOsm. For measurement of resurgent current, 200 μM β4 peptide was included in the internal solution. Recordings were acquired with an Axopatch 200B amplifier (Molecular Devices, Sunnyvale, CA) at 50 kHz and filtered at 5 kHz. Leak subtraction was performed with a P/4 protocol. Fast and slow capacitive currents were compensated. After achieving the whole-cell configuration, recordings began after 10 minutes to allow for equilibration of cell contents with the internal solution. Series resistance was compensated 80-90%, and cells with unstable series resistance or series resistance >8 MOhms were discarded from further analysis.

Protocols for activation, inactivation, recovery from inactivation, and resurgent current are depicted in the figures. To measure residual current after high-frequency pulses, cells were held at −120 mV and depolarized to 0 mV for 5 ms for 300 pulses at the indicated frequency. The residual current was calculated as the ratio of the 300^th^ pulse to the 1^st^ pulse. Resurgent current was assessed by first depolarizing the cell to +30 mV, followed by repolarization steps at the indicated voltages. Persistent current was calculated as the average current of the last 10 ms of a 200 ms depolarizing pulse to 0 mV, normalized to the peak current from a TTX-subtracted record.

### Data Analysis

Analysis of electrophysiological data was performed with ClampFit (Molecular Devices), Microsoft Excel, MatLab (Mathworks, Natick, MA), and GraphPad Prism (San Diego, CA). All data points are represented as mean ± SEM, and *n* is the sample size. Current densities were calculated by dividing peak currents at each potential by the membrane capacitance obtained during recording. Conductance was calculated using the equation G = I*(V - E_rev_), where E_rev_ is the calculated sodium reversal potential. Activation and inactivation V_1/2_ values were obtained by fitting the normalized conductance (activation) or the normalized current (inactivation) to a Boltzmann equation: 1/(1+exp[(V - V_1/2_)/k]), where V_1/2_ is the half-activation voltage and k is the slope factor. Inactivation time-course was estimated by fitting a single-component exponential curve to the decay portion of the sodium current using ClampFit. Recovery from inactivation was fit with a biexponential curve using Prism. The data distribution was assessed for normality with a Shapiro-Wilk test. If the data were normally distributed, a parametric test (t-test) was used to assess significance; if not, a nonparametric test was used (Mann-Whitney U-test). Statistical significance was set at P<0.05. When assessing three or more groups (as in the carbamazepine experiment), the data were analyzed with a repeated measures ANOVA with Dunnett’s multiple comparisons post-hoc test, comparing each drug response to the control condition.

### Reagents

Tetrodotoxin (TTX; Tocris Bioscience, Minneapolis, MN) was prepared as a 1 mM stock solution in water, then diluted to 500 nM in the external solution. Carbamazepine (Sigma) was prepared as a 250 mM stock solution in DMSO and diluted to the indicated concentration in the external solution. A β4 peptide (KKLITFILKKTREK-OH), generously supplied by Dr. Chris Ahern at the University of Iowa, was diluted in the internal solution to a final concentration of 200 μM and used immediately or frozen in aliquots.

### Molecular Modeling

A structure of the C-Terminal domain of human Na_V_1.2 determined by X-ray crystallography at 3.02 Angstrom resolution in complex with calmodulin and FGF13 was obtained from chain B of PDB code 4JPZ.^12^ The Met-1879 residue was changed to Thr *in silico* using UCSF Chimera (RBVI, San Francisco, CA), with the highest probability rotamer selected. This M1879T structure and the original structure (WT) were then energy minimized using 1000 steepest descent and 10 conjugate gradient steps. Contacts of non-backbone residues with either Met-1879 or Thr-1879 were identified using Chimera with default parameters.

## RESULTS

### Case Report

An infant boy was evaluated for a seizure disorder accompanied by mild global developmental delay with onset at age two months. He had a normal birth history, and no family history of seizures or epilepsy. The child initially came to medical attention for an apneic event, which was diagnosed as reflux. EEG and brain MRI were performed which were normal. Events recurred and he was referred to Lurie Children’s and diagnosed with tonic seizures accompanied by apneic spells and cyanosis. Neurological exam was notable for low axial and appendicular muscle tone. He was treated with levetiracetam, pyridoxine, and topiramate, yet continued to have daily seizures. Following each acute loading dose of intravenous fosphenytoin he became seizure free for two days, but then seizures recurred. Transition to oral phenytoin maintained seizure freedom. Genetic testing revealed a *de novo SCN2A* variant (c.5636T>C) predicting a novel missense mutation (p.Met1879Thr, M1879T) in the Na_V_1.2 voltage-gated sodium channel. On follow-up, there had been difficulty maintaining adequate phenytoin levels, and he was transitioned to carbamazepine monotherapy with continued seizure control. At last follow up after 2 years old, he has developmental delay and hyperactivity with autistic features. He had minimal receptive and expressive language, can feed himself, and can run with an ataxic gait.

### Functional consequences of SCN2A-M1879T

We investigated the functional effect of the M1879T mutation using recombinant human Na_V_1.2 expressed in a heterologous cell line (HEK293T) stably expressing the human β1 and β2 subunits. Representative voltage-dependent currents recorded from WT and M1879T expressing cells are depicted in **Figure 1A** and summary data for all functional properties are presented in **Table 1**. The peak current density was not significantly different between WT and M1879T channels (**Figure 1B**). However, the time-course of inactivation was significantly slower for M1879T compared to WT channels (**Figure 1C**, upper). A voltage ramp protocol evoked a pronounced aberrant current in cells expressing M1879T but not in WT expressing cells (**Figure 1C,** lower). Inactivation time constants reveal a significant impairment of M1879T inactivation compared to WT across the range of tested voltages (**Figure 1D**). M1879T mutant channels did not exhibit a significant difference in persistent sodium current measured as a percentage of peak current. These findings are consistent with impaired fast inactivation for the mutant channel that predicts greater sodium conductance over time following channel activation.

**Figure 1.**
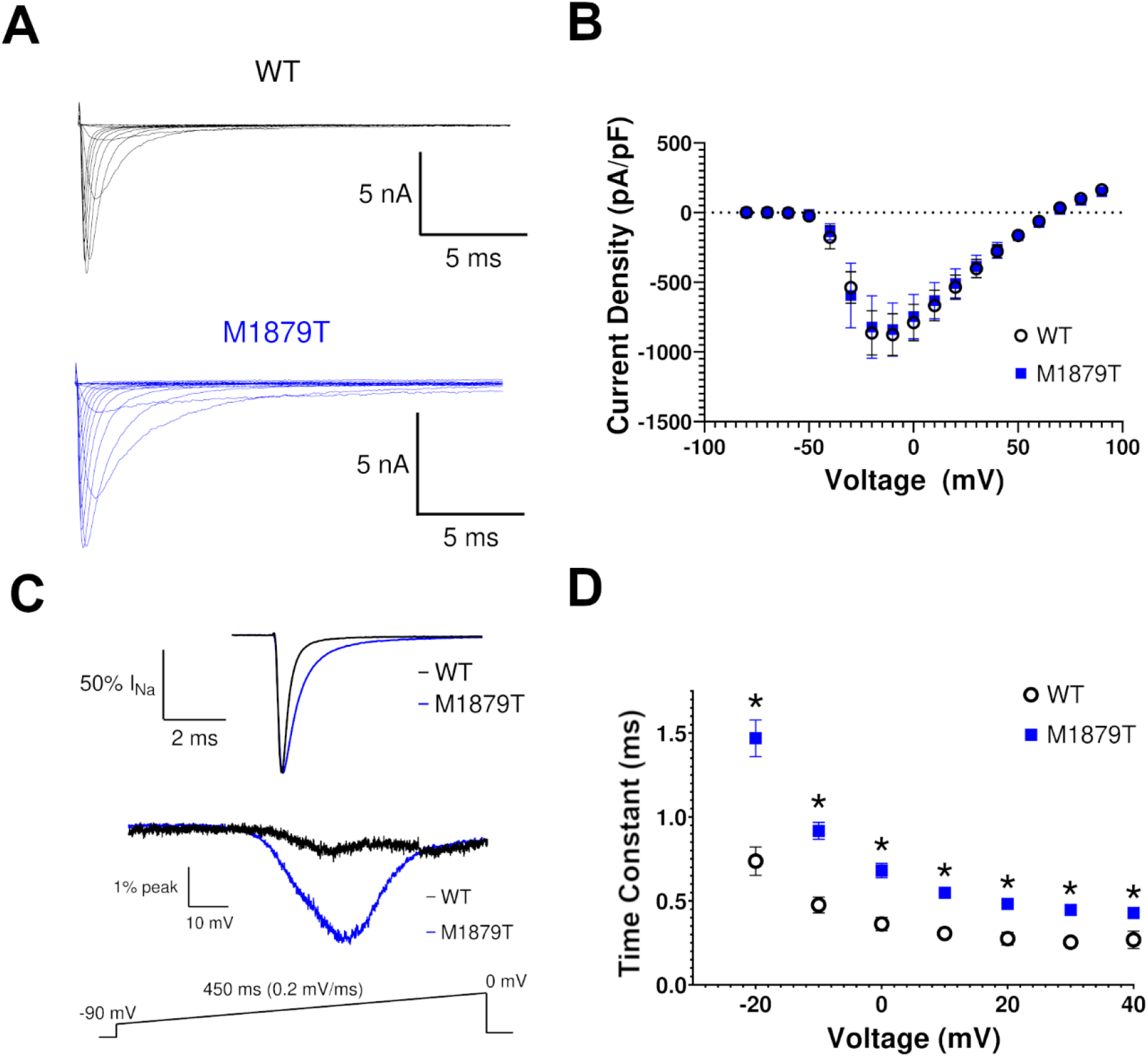
M1879T mutation alters Na_V_1.2 inactivation kinetics. (A) Representative current recordings from WT and M1879T Na_V_1.2 obtained using the voltage protocol shown in Figure 2A (inset). (B) Current density-voltage plot for WT (n=14) and M1879T (n=14) channels. (C) Top, average of six TTX-subtracted recorded at 0 mV and normalized to peak current to highlight inactivation time-course of WT (black) and M1879T (blue). Bottom, average currents for WT (black, average of 5) and M1879T (blue, average of 6) elicited using the voltage ramp protocol shown below. Traces were normalized to peak current measured at 0 mV. (D) Voltagedependence of inactivation time constants determined from single-component exponential curve fits to the current decay. n=14 for WT and n=14 for M1879T, *p<0.05 by Mann-Whitney U test.

**Table 1.** Summary of functional properties for WT and M1879T channels

|  | **WT (n)** | **M1879T (n)** | **M1879T-neonatal (n)** |
| --- | --- | --- | --- |
| Peak current density (pA/pF) | -972 ± 35 (14) | -908 ± 163 (14) | -654 ± 125 (12) |
| Inactivation time constant (τ, ms) 0 mV | 0.36 ± 0.04 (14) | 0.68 ± 0.04* (14) | 0.69 ± 0.05 (12) |
| Activation voltage-dependence |  |  |  |
| V<sub>1/2</sub> (mV) | -29.2 ± 1.7 (14) | -28.2 ± 1.8 (14) | -27.7 ± 1.2 (14) |
| Slope factor (k) | 4.8 ± 0.5 (14) | 4.5 ± 0.4 (14) | 4.8 ± 0.4 (14) |
| Inactivation voltage-dependence |  |  |  |
| V<sub>1/2</sub> (mV) | -66.3 ± 1.1 (18) | -53.8 ± 1.1* (25) | -50.3 ± 1.4 (15) |
| Slope factor (k) | -6.1 ± 0.2 (18) | -7.4 ± 0.3* (25) | -7.8 ± 0.3 (15) |
| Recovery from inactivation |  |  |  |
| τ<sub>slow</sub> (ms) | 154 ± 88 (10) | 139 ± 35 (10) | 132 ± 28 (11) |
| τ<sub>fast</sub> (ms) | 2.18 ± 0.30 (10) | 2.48 ± 0.32 (10) | 2.96 ± 0.44 (11) |
| % fast component | 77.7 ± 1.8 (10) | 76.7 ± 4.4 (10) | 78.3 ± 3.1 (11) |
| Persistent Current (% of peak) | 0.33 ± 0.05 (9) | 0.21 ± 0.04 (9) | 0.47 ± 0.11 (8) |

We compared the voltage-dependence of channel activation and steady-state inactivation between M1879T and WT channels. The voltage dependence of activation was not different between M1879T and WT channels (**Figure 2A**). By contrast, cells expressing M1879T exhibited a significantly depolarized steady-state inactivation curve compared to WT expressing cells (**Figure 2B**). These results are also consistent with altered fast inactivation and predict greater sodium conductance by mutant channels evoked by membrane depolarizations within the physiological range.

**Figure 2.**
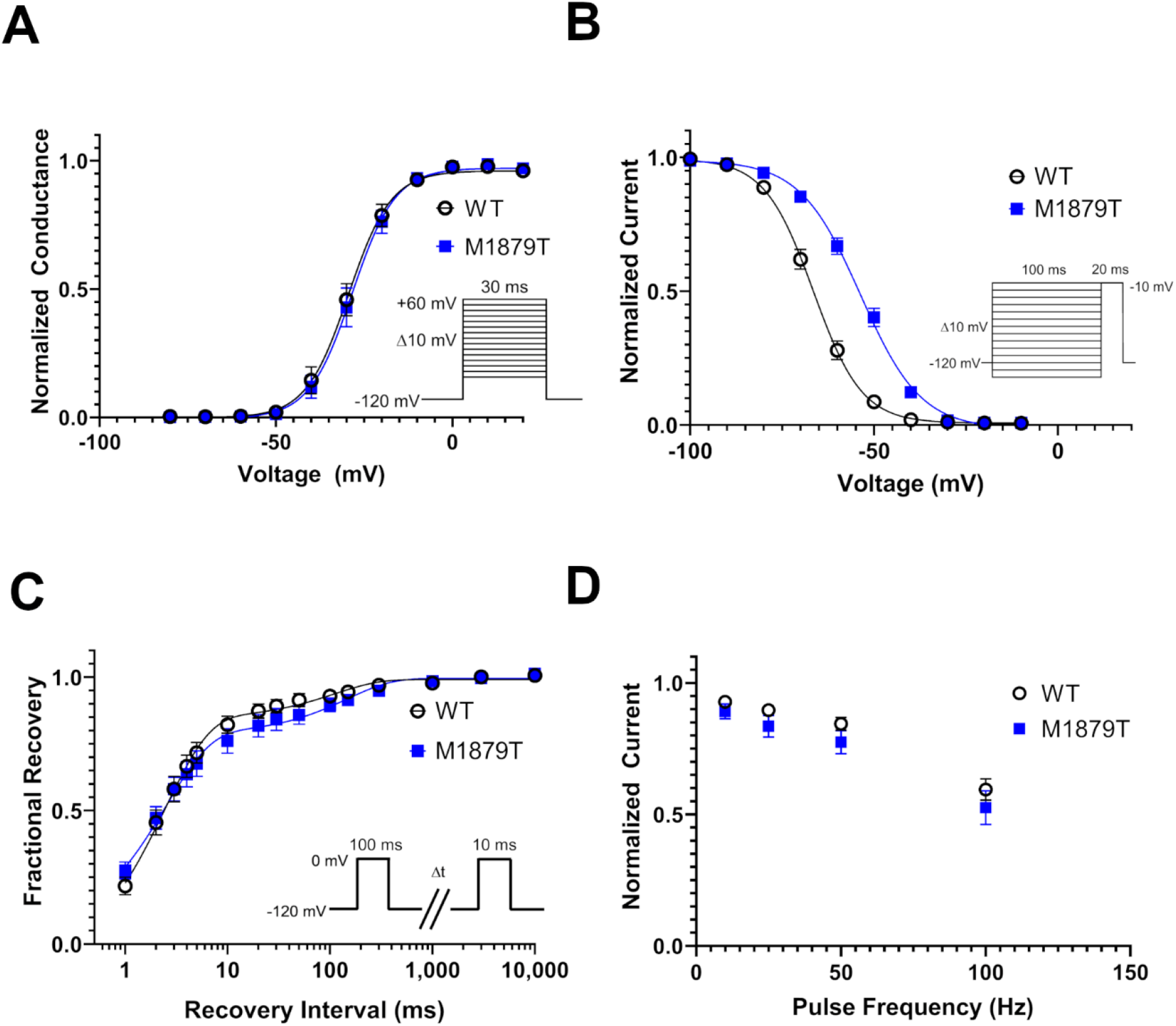
M1879T affects voltage-dependence of inactivation. (A) Voltage-dependence of activation of WT (n=14) and M1879T (n=14) channels determined using the voltage protocol shown as an inset. There was no difference in activation V_1/2_ between WT and M1879T (p=0.51 by Mann-Whitney U test). (B) Voltage-dependence of inactivation of WT (n=18) and M1879T (n=25) channels (voltage protocol shown as an inset). There was a significant depolarizing shift in inactivation V_1/2_ (p<0.0001 by Mann-Whitney U test). (C) Time course of recovery from inactivation after 100 ms depolarization (protocol shown as inset) comparing WT (n=10) and M1879T (n=10). (D) Plot of residual current (comparing 300^th^ pulse to 1^st^ pulse) after repetitive pulsing to 0 mV at the indicated frequency (n=7 for WT and n=7 for M1879T). There were no significant differences between WT and M1879T at any frequency (p>0.05 by Mann-Whitney U test).

Recovery from inactivation, assessed using a two-pulse protocol, followed a biexponential time course and was not significantly different between M1879T and WT channels (**Figure 2C**). Similarly, sodium channel availability during depolarizing pulse trains at different frequencies (10, 25, 50, and 100 Hz) were not significantly different between mutant and WT channels (**Figure 2D**). These results suggest that M1879T selectively affects the kinetics of onset and voltage-dependence of channel inactivation.

*SCN2A* undergoes developmentally regulated alternative mRNA splicing that leads to incorporation of an alternate exon encoding a portion of the domain I voltage-sensor domain (S3 and S4 helices). Because *SCN2A* variants associated with early onset epilepsy may exhibit more severe functional consequences in the splice variant expressed preferentially during early development,^13^ we tested whether the substitution of the neonatal exon would affect the properties of the M1879T mutant channel. In this case, none of the key functional properties (**Supplemental Figure S1**) were significantly different between the adult and neonatal isoforms of the M1879T channel.

### M1879T channels exhibit enhanced resurgent current

We next compared the level of resurgent current between WT and M1879T channels. These experiments were motivated by a prior report indicating that the epilepsy-associated mutation *SCN2A*-R1882Q affecting a nearby residue had greater resurgent current than WT channels.^14^ We recorded resurgent current in WT and M1879T expressing cells elicited by the voltage protocol shown in **Figure 3A** (top) with the β4 peptide in the internal solution. M1879T channels (**Figure 3A**, bottom) exhibited larger resurgent currents than WT channels (**Figure 3A**, middle). This difference was evident across several tested voltages with comparisons made using either current density (**Figure 3B**) or the level of resurgent current measured as a percentage of the peak current elicited at −10 mV (**Figure 3C**). Enhanced resurgent current exhibited by M1879T channels may contribute to the gain of sodium conductance conferred by this mutation.

**Figure 3.**
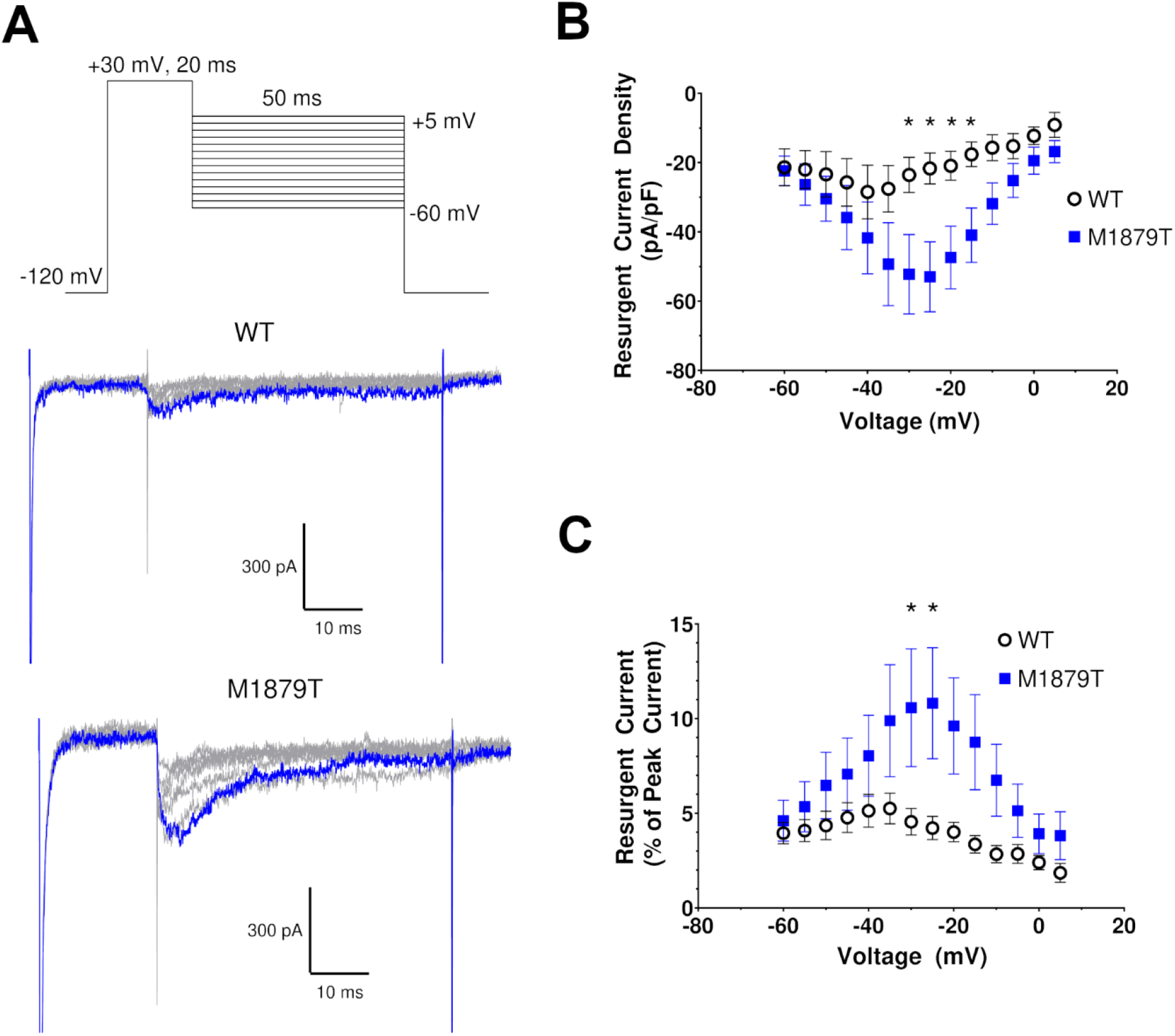
M1879T exhibits enhanced resurgent current. (A) Top, voltage protocol for eliciting resurgent current. Middle (WT) and bottom (M1879T) panels show representative current traces using protocol above (10 mV voltage steps are illustrated). The blue trace represents the peak resurgent current elicited at −20 mV. (B) Voltage dependence of resurgent current density for WT and M1879T (*, p<0.05 by Mann-Whitney U test). (C) Plot of resurgent current expressed as % of peak transient current for WT and M1879T (*, p<0.05 by Mann-Whitney U test).

### Effect of carbamazepine on M1879T channels

Because carbamazepine (CBZ) was effective as monotherapy in the reported case, we examined the effects of this drug on M1879T channels (adult splice isoform). As shown in **Figure 4A**, application of 100 μM and 300 μM CBZ caused a dose-dependent hyperpolarizing shift in steady-state inactivation (control [no drug]: V_1/2_ = −48.5; 100 μM CBZ: V_1/2_ = −52.0; 300 μM CBZ: V_1/2_ = −55.8; n=6). By comparing each treated cell to its own control value, we determined that 100 μM and 300 μM CBZ caused 3.5 ± 0.7 mV and 8.5 ± 0.7 mV hyperpolarizing shifts in steady-state inactivation V_1/2_, respectively (**Figure 4B**). Vehicle (DMSO) treatment did not cause a significant shift in steady-state inactivation (data not shown). Furthermore, CBZ reduced the aberrant ramp current exhibited by M1879T channels (**Figure 4C**). Finally, CBZ was associated with a significantly lower degree of channel availability after high-frequency pulses at 100 Hz (fractional availability after 300^th^ pulse: control: 0.61; 100 μM CBZ: 0.49; 300 μM CBZ: 0.32, n=5; p=0.0105 for control vs.100 μM CBZ and p=0.0014 for control vs. 300 μM CBZ), but not at lower pulsing frequencies (**Figure 4D**). The partial normalization of voltage-dependence of inactivation coupled with use-dependent block of the mutant channel correlate with the clinical efficacy of CBZ in the reported case.

**Figure 4.**
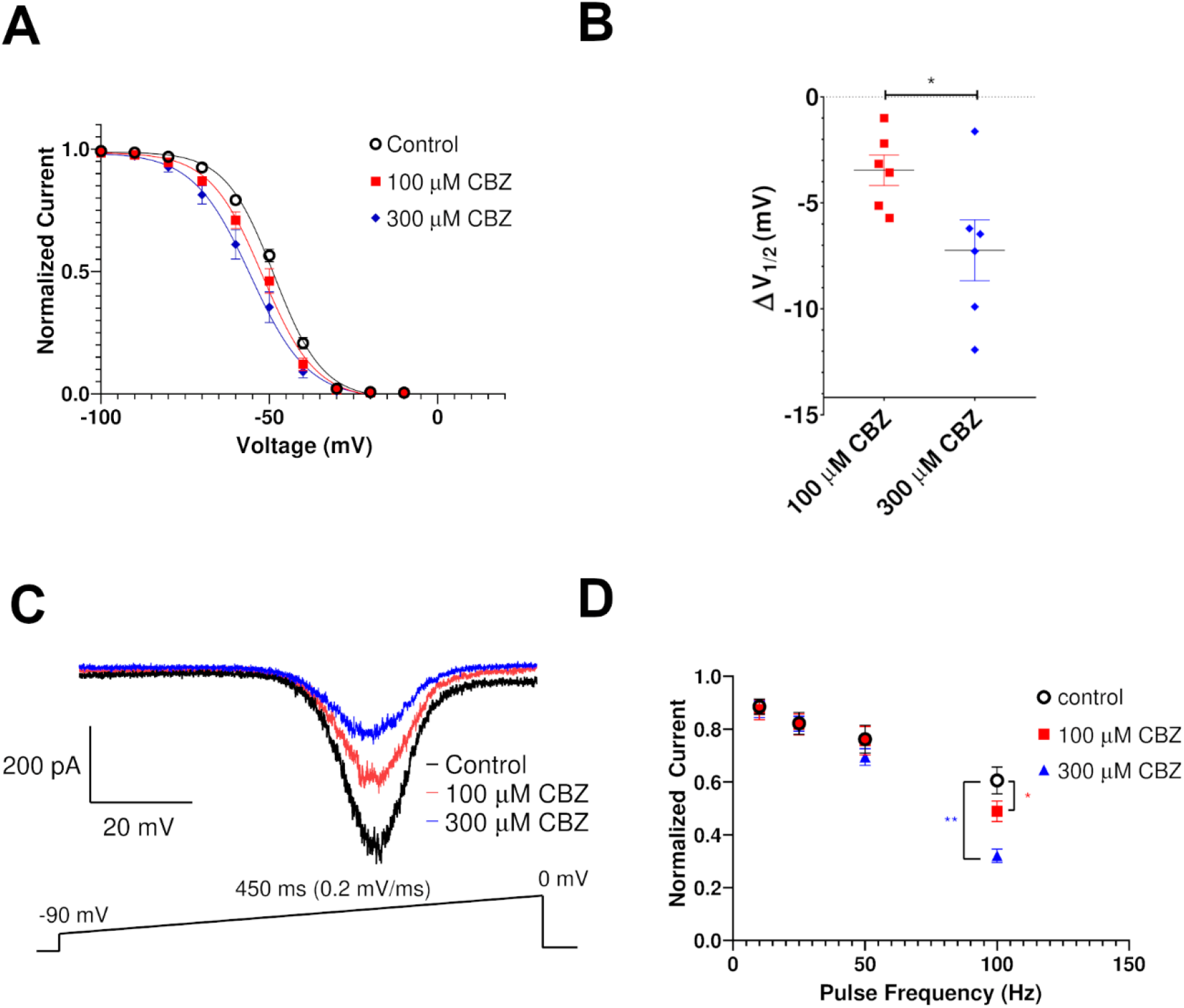
Effects of carbamazepine on M1879T channels. (A) Carbamazepine (CBZ) induces a hyperpolarizing shift of the voltage-dependence of inactivation (n=6). (B) Plot of change in inactivation V_1/2_ in response to 100 μM and 300 μM carbamazepine (n=6; *, p<0.05 by paired t-test). (C) Ramp current determined in the absence (control) or presence of 100 μM CBZ (red trace) and 300 μM CBZ (blue trace). (D) Plot of residual current (comparing 300^th^ pulse to 1^st^ pulse) after 0 mV pulses at the indicated frequency. At 100 Hz, there was a significant difference in residual current in the presence of 100 μM and 300 μM CBZ compared to control conditions (one-way ANOVA with repeated measures and Dunnett’s post-hoc test, *p < 0.05 for 100 μM vs. control; **p < 0.05 for 300 μM vs. CBZ; n=5).

We also assessed whether CBZ treatment affects resurgent current. Cells expressing M1879T were treated with either 300 μM CBZ or DMSO vehicle. Cells treated with DMSO only (**Figure 5A, left**) exhibited a time-dependent rise in resurgent current, likely related to the time-dependent diffusion of the β4 peptide into the intracellular compartment seen in control solution (data not shown). In contrast, cells treated with CBZ showed lower resurgent current levels compared to the control condition (**Figure 5A, B**). Specifically, DMSO-treated cells exhibited a 22.1 ± 4.6% (n=7) rise in maximal resurgent current density compared to control, while CBZ-treated cells had a 21.2 ± 3.1% (n=7) fall compared to the control condition (**Figure 5C**). Because CBZ affects peak current, we considered the possibility that reduced resurgent current merely reflects a proportional reduction in the total current. Indeed, 300 μM CBZ treatment caused a significant change in peak current compared to DMSO treatment (**Figure 6A**). When this difference was taken into account by examining the maximal resurgent current as a percentage of the peak current, there remained a significant difference between the DMSO-treated and CBZ-treated cells (**Figure 6B**). Taken together, these data indicate that 300 μM CBZ likely attenuates resurgent current independent of its effect on peak current, but this effect is modest.

**Figure 5.**
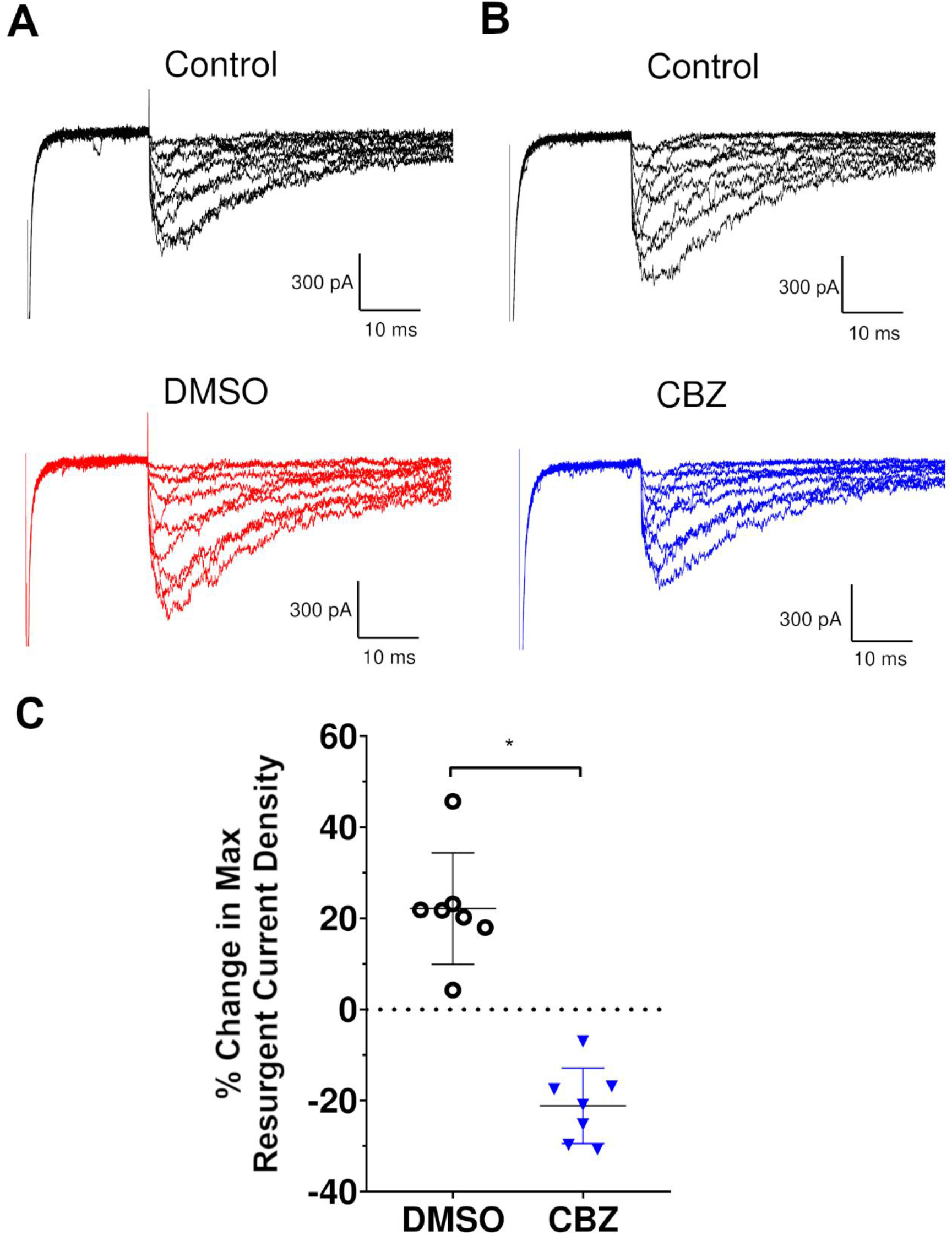
Effect of carbamazepine on resurgent current. (A) Representative M1879T resurgent current traces before (top) and after DMSO (bottom) treatment. (B) Representative M1879T resurgent current before (top) and after CBZ (bottom) treatment. (C) Percent change in maximum resurgent current density, comparing DMSO and CBZ (n = 7; *, p<0.0001 by unpaired t test).

**Figure 6.**
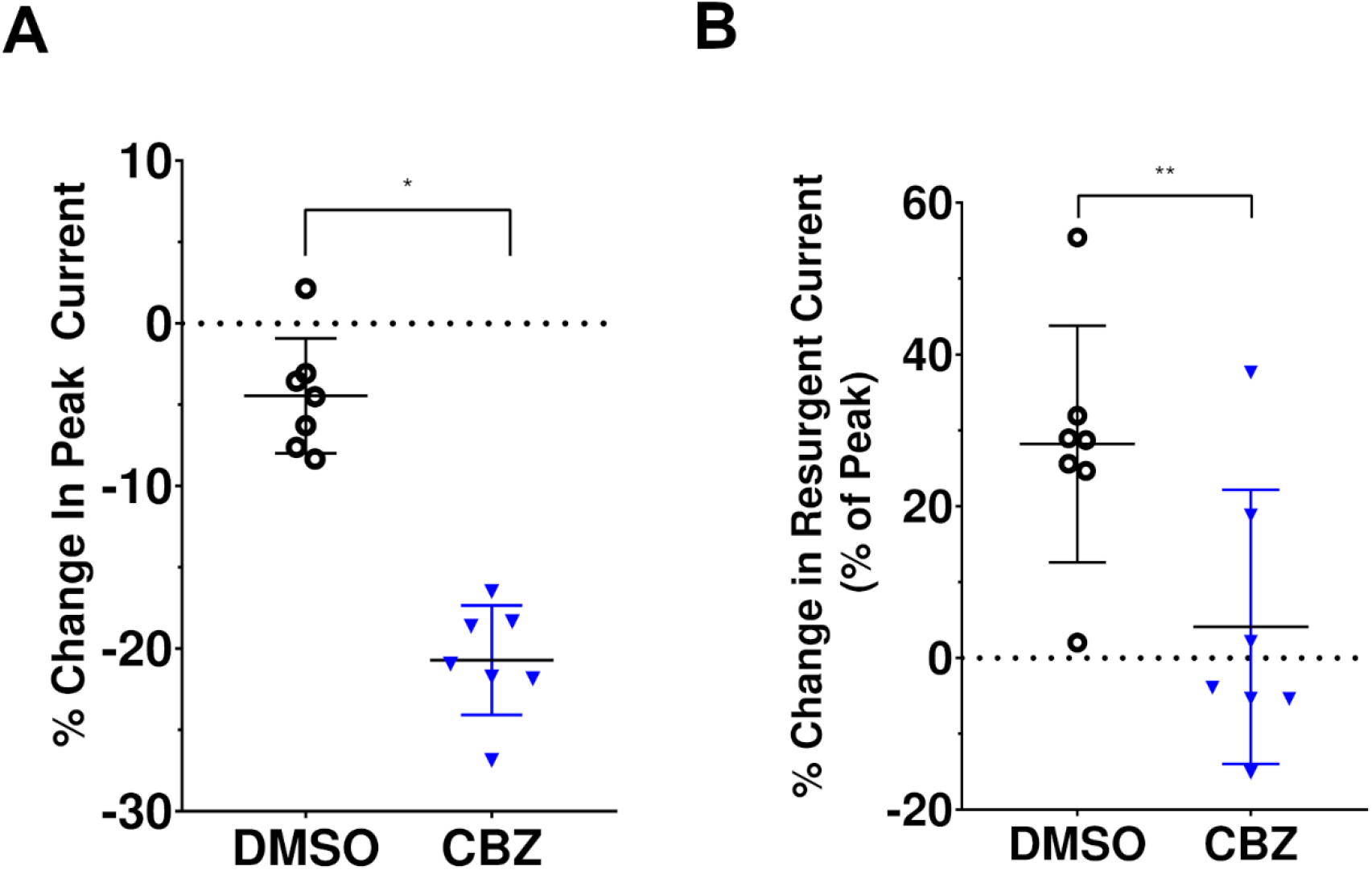
Comparison of carbamazepine effects on M1879T peak transient and resurgent current. (A) Percent change in peak transient current (DMSO: −4.5 ± 1.3% vs. CBZ: −20.7 ± 1.3%; n=7 both groups; *, p<0.0001 by unpaired t test). (B) Percent change in resurgent current (as percent of peak transient current; DMSO: 28.2 ± 5.9% vs. CBZ: 4.1 ± 6.8% of peak, n=7 for both groups; **, p=0.02 by unpaired t test).

## DISCUSSION

Our study investigated a *de novo SCN2A* coding sequence variant (p.M1879T) associated with infantile-onset epilepsy. The child we report exhibited frequent tonic seizures complicated by apnea unresponsive to levetiracetam and topiramate then subsequently had a dramatic response to the sodium channel blockers phenytoin and later carbamazepine. Initially the variant was categorized as a variant of uncertain significance (VUS), but electrophysiological evaluation of the variant sodium channel revealed severely impaired inactivation gating consistent with a gain-of-function. Specifically, we observed a substantially depolarized voltage-dependence of inactivation, slower time course of inactivation, and enhanced resurgent current. Unlike other *SCN2A* variants associated with early-onset epilepsy, expression of the variant in the neonatal splice variant of the channel was not associated with greater dysfunction.^13^ Carbamazepine partially normalized the voltage-dependent inactivation of the mutant channel, dampened resurgent current, and exhibited potent frequency-dependent block of the mutant channel. Therefore, our study demonstrates concordance of *in vitro* pharmacology using heterologous cells with the drug response observed clinically for the affected child.

The novel missense mutation we report in this paper affects residue Met-1879 located in the distal portion of the intracellular carboxyl-terminal domain (CTD), upstream of a conserved IQ domain necessary for calmodulin binding.^15–17^ This methionine residue is absolutely conserved in the mammalian voltage-gated sodium channel family. Variants affecting orthologous sodium channels predicting the same methionine to threonine amino acid substitution occur at corresponding positions in the cardiac sodium channel (Na_V_1.5, M1875T) associated with familial atrial fibrillation,^18^ and in a peripheral nerve sodium channel (Na_V_1.7, M1870T) associated with painful diabetic peripheral neuropathy.^19^ Both orthologous mutations exhibited similar biophysical effects as *SCN2A*-M1879T in heterologous cells including a depolarizing shift in the voltage-dependent inactivation and slowed inactivation kinetics without affecting voltage-dependent activation or persistent current. These results along with our findings implicate Met-1879 as important for inactivation properties, and mutation to threonine appears to destabilize the inactivated state of the channel.

Other epilepsy-associated mutations have been identified in the Na_V_1.2 CTD most notably R1882Q, which causes early-onset epilepsy of varying severity and drug-responsiveness. The R1882Q channel exhibits a depolarized voltage-dependence of inactivation, as well as an increase in non-inactivating persistent current compared to wild-type channels, an attribute we did not observe for the M1879T channel. This combination of functional anomalies is predicted to promote greater neuronal excitability.^20^

The molecular basis for channel dysfunction associated with the M1879T mutation can be considered in the context of recent advances in the structural biology of sodium channels. In the past few years, there has been an explosion of available structures for eukaryotic sodium channels, including human Na_V_1.2.^21^ Unfortunately, the intracellular regions including the CTD are poorly resolved or missing in most of these structures except in the full-length sodium channel from the American cockroach (Na_V_Pas).^22^ Fortunately, there are structural data available for the isolated Na_V_1.2 CTD in complex with calmodulin and FGF13.^12^ In this structure, Met-1879 resides within an alpha-helix that forms a helix-loop-helix motif neighboring the IQ domain needed for calmodulin binding. The proximity of this residue to the base of the fourth voltage-sensing domain provides a plausible explanation for how the M1879T mutation might affect inactivation driven by this domain.^23^ In this structure, the side chain of Met-1879 projects outward to a nearby helix in the CTD where it forms contacts with a network of side chains belonging to nearby residues (**Figure 7A**). Molecular modeling suggests that the M1879T variant, with its smaller side chain, eliminates several of these intramolecular contacts that may stabilize inactivation (**Figure 7B**).

**Fig. 7.**
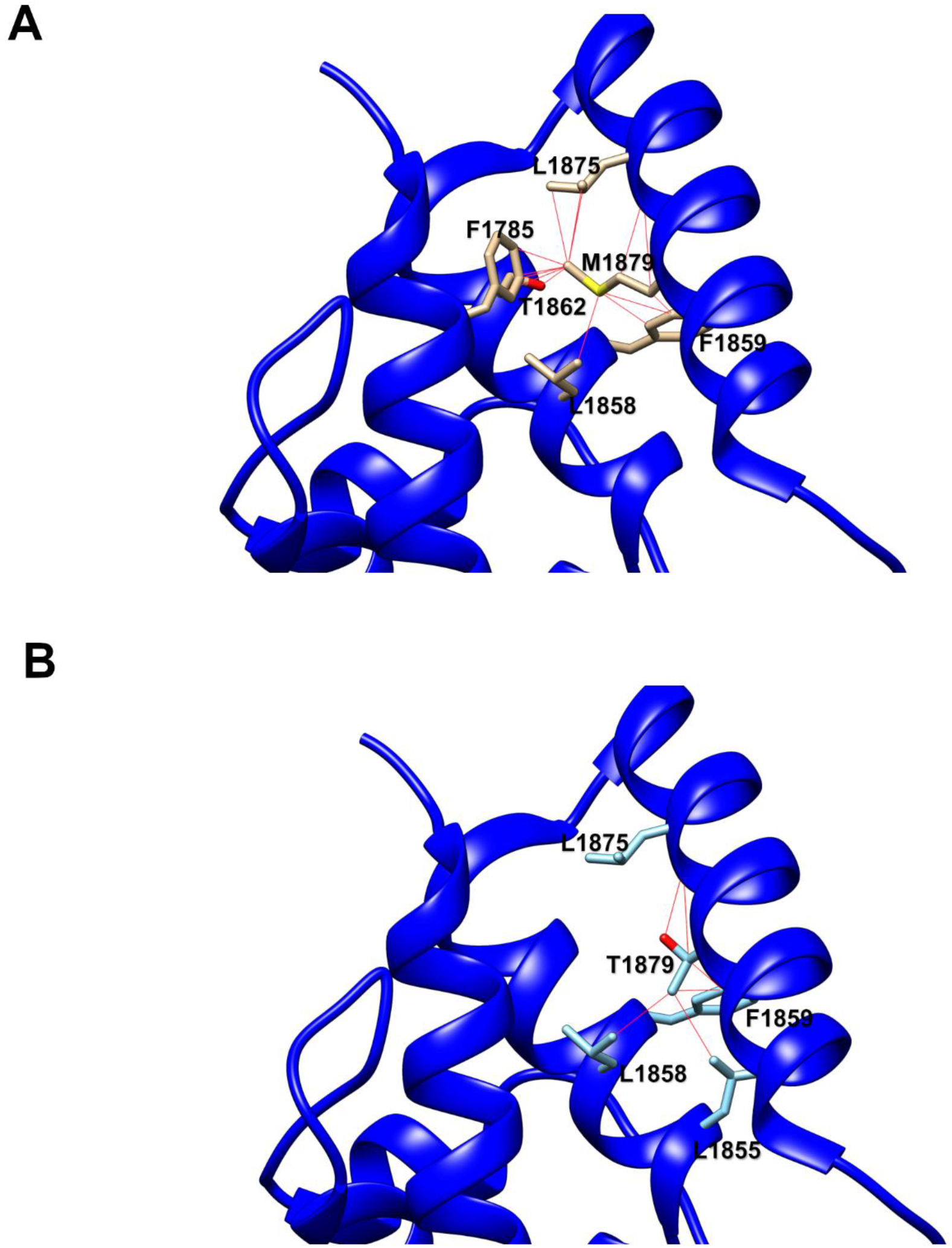
Molecular modeling of Met-1879 and Thr-1879 contacts in Na_V_1.2 C-terminal domain. (A) Ribbon diagram of WT channel showing specific contacts between Met-1879 and interacting residues (red lines). (B) Ribbon diagram of M1879T channel showing specific contacts between Thr-1879 and interacting residues (red lines).

An understudied functional effect of *SCN2A* mutations is resurgent current, and only two variants (R853Q, R1882Q) have been investigated previously for this phenomenon in heterologous cells.^14^ R853Q channels exhibit less resurgent current compared to WT channels, whereas R1882Q has a much larger resurgent current compared, similar to what we observed with M1879T. However, the impact of resurgent current on promoting epileptic activity is unclear. Resurgent currents have been demonstrated for other sodium channelopathies, including those implicated in pain, myotonia, long-QT syndrome, and *SCN8A*-related epilepsy.^24–26^

We examined the effect of carbamazepine on resurgent current to determine if suppression of this anomalous channel activity might explain the antiepileptic effects observed in this patient. Although high CBZ concentration (300 μM) did significantly suppress resurgent current, the reduction we observed correlated with an overall reduction in peak current. When resurgent current was normalized to peak current, the effect of carbamazepine appeared modest. These experimental findings suggested that the principal effect of carbamazepine on reducing seizures associated with the M1879T variant more likely relates to changes in voltage-dependent inactivation and channel inhibition at high-frequency stimulation rather than attenuation of resurgent current.

Suppression of resurgent current by other agents has also been demonstrated. The endogenous cannabinoid anandamide inhibits resurgent current without affecting transient current,^27^ whereas PRX-330, a novel anti-epileptic drug candidate, inhibits both persistent and resurgent currents in a mouse model of SCN8A encephalopathy.^28^ Cannabidiol (CBD), a recently approved anti-epileptic drug for the treatment of Dravet and Lennox-Gastaut syndromes, also inhibits resurgent current more than transient current associated with two epilepsy-associated *SCN8A* variants (L1331V, N1768D).^25^ Further investigations of resurgent current pharmacology are warranted.

In summary, our study illustrates that functional and pharmacological investigations of *SCN2A* variants in heterologous cells has value in defining genotype-phenotype correlations, demonstrating molecular mechanisms of disease and drug responses, and contributing to our knowledge of structure-functional relationships for the broader voltage-gated sodium channel family.

## ACKNOWLEDGEMENTS

The authors sincerely thank the patient and his family for their inclusion in this study. This work was supported by National Institute of Neurological Disorders and Stroke grants R25-NS070695 (S.K.A.) and U54-NS108874 (A.L.G.), and a generous gift from the Davee Foundation (A.L.G.).

## AUTHORS’ CONTRIBUTIONS

The study was conceived by SKA, JMM and ALG. Experiments were designed by SKA and ALG with guidance from CHT. Experiments were performed and data analyzed by SKA. JMD, CHT, and TA performed mutagenesis and development of cell lines. SKA and ALG wrote the manuscript with input from all co-authors.

## REFERENCES

1. Brunklaus A, Du J, Steckler F, et al. Biological concepts in human sodium channel epilepsies and their relevance in clinical practice. Epilepsia 2020;DOI:10.1111/epi.16438

2. Reynolds C, King MD, Gorman KM. The phenotypic spectrum of SCN2A-related epilepsy. Eur. J. Paediatr. Neurol. 2020;24:117–122;DOI:10.1016/j.ejpn.2019.12.016

3. Brunklaus A. Precision medicine in sodium channelopathies – Moving beyond seizure control towards disease modification. Eur. J. Paediatr. Neurol. 2020;24:7;DOI:10.1016/j.ejpn.2020.01.008

4. Ben-Shalom R, Keeshen CM, Berrios KN, et al. Opposing Effects on Na_V_1.2 Function Underlie Differences Between SCN2A Variants Observed in Individuals With Autism Spectrum Disorder or Infantile Seizures. Biol. Psychiatry 2017;82(3):224–232;DOI:10.1016/j.biopsych.2017.01.009

5. Symonds JD, McTague A. Epilepsy and developmental disorders: Next generation sequencing in the clinic. Eur. J. Paediatr. Neurol. 2020;24:15–23;DOI:10.1016/j.ejpn.2019.12.008

6. Hu W, Tian C, Li T, et al. Distinct contributions of Na_v_1.6 and Na_v_1.2 in action potential initiation and backpropagation. Nat. Neurosci. 2009;12(8):996–1002;DOI:10.1038/nn.2359

7. Tian C, Wang K, Ke W, et al. Molecular identity of axonal sodium channels in human cortical pyramidal cells. Front. Cell. Neurosci. 2014;8;DOI:10.3389/fncel.2014.00297

8. Liao Y, Deprez L, Maljevic S, et al. Molecular correlates of age-dependent seizures in an inherited neonatal-infantile epilepsy. Brain 2010;133(Pt 5):1403–14;DOI:10.1093/brain/awq057

9. Sanders SJ, Campbell AJ, Cottrell JR, et al. Progress in Understanding and Treating SCN2A-Mediated Disorders. Trends Neurosci. 2018;41(7):442–456;DOI:10.1016/j.tins.2018.03.011

10. Musto E, Gardella E, Møller RS. Recent advances in treatment of epilepsy-related sodium channelopathies. Eur. J. Paediatr. Neurol. 2020;24:123–128;DOI:10.1016/j.ejpn.2019.12.009

11. Kahlig KM, Saridey SK, Kaja A, et al. Multiplexed transposon-mediated stable gene transfer in human cells. Proc. Natl. Acad. Sci. U. S. A. 2010;107(4):1343–1348;DOI:10.1073/pnas.0910383107

12. Wang C, Chung BC, Yan H, et al. Structural analyses of Ca^2+^/CaM interaction with Na_V_ channel C-termini reveal mechanisms of calcium-dependent regulation. Nat. Commun. 2014;5:4896;DOI:10.1038/ncomms5896

13. Thompson CH, Ben-Shalom R, Bender KJ, George AL. Alternative splicing potentiates dysfunction of early-onset epileptic encephalopathy SCN2A variants. J. Gen. Physiol. 2020;152(3);DOI:10.1085/jgp.201912442

14. Mason ER, Wu F, Patel RR, et al. Resurgent and gating pore currents induced by De Novo SCN2A epilepsy mutations. eNeuro 2019;6(5);DOI:10.1523/ENEURO.0141-19.2019

15. Mori M, Konno T, Ozawa T, et al. Novel interaction of the voltage-dependent sodium channel (VDSC) with calmodulin: Does VDSC acquire calmodulin-mediated Ca^2+^ - sensitivity? Biochemistry 2000;39(6):1316–1323;DOI:10.1021/bi9912600

16. Kim J, Ghosh S, Liu H, et al. Calmodulin mediates Ca^2+^ sensitivity of sodium channels. J. Biol. Chem. 2004;279(43):45004–45012;DOI:10.1074/jbc.M407286200

17. Theoharis NT, Sorensen BR, Theisen-Toupal J, Shea MA. The neuronal voltage-dependent sodium channel type II IQ motif lowers the calcium affinity of the C-domain of calmodulin. Biochemistry 2008;47(1):112–123;DOI:10.1021/bi7013129

18. Makiyama T, Akao M, Shizuta S, et al. A Novel SCN5A Gain-of-Function Mutation M1875T Associated With Familial Atrial Fibrillation. J. Am. Coll. Cardiol. 2008;52(16):1326–1334;DOI:10.1016/j.jacc.2008.07.013

19. Blesneac I, Themistocleous AC, Fratter C, et al. Rare Na_V_1.7 variants associated with painful diabetic peripheral neuropathy. Pain 2018;159(3):469–480;DOI:10.1097/j.pain.0000000000001116

20. Berecki G, Howell KB, Deerasooriya YH, et al. Dynamic action potential clamp predicts functional separation in mild familial and severe de novo forms of SCN2A epilepsy. Proc. Natl. Acad. Sci. U. S. A. 2018;115(24):E5516–E5525;DOI:10.1073/pnas.1800077115

21. Pan X, Li Z, Huang X, et al. Molecular basis for pore blockade of human Na^+^ channel Na_v_1.2 by the μ-conotoxin KIIIA. Science 2019;363(6433):1309–1313;DOI:10.1126/science.aaw2999

22. Shen H, Zhou Q, Pan X, et al. Structure of a eukaryotic voltage-gated sodium channel at near-atomic resolution. Science (80-.). 2017;355(6328);DOI:10.1126/science.aal4326

23. Capes DL, Goldschen-Ohm MP, Arcisio-Miranda M, et al. Domain IV voltage-sensor movement is both sufficient and rate limiting for fast inactivation in sodium channels. J. Gen. Physiol. 2013;142(2):101–112;DOI:10.1085/jgp.201310998

24. Jarecki BW, Piekarz AD, Jackson JO, Cummins TR. Human voltage-gated sodium channel mutations that cause inherited neuronal and muscle channelopathies increase resurgent sodium currents. J. Clin. Invest. 2010;120(1):369–378;DOI:10.1172/JCI40801

25. Patel RR, Barbosa C, Brustovetsky T, et al. Aberrant epilepsy-associated mutant Na_v_1.6 sodium channel activity can be targeted with cannabidiol. Brain 2016;139(8):2164–2181;DOI:10.1093/brain/aww129

26. Xiao Y, Barbosa C, Pei Z, et al. Increased Resurgent Sodium Currents in Na_v_1.8 Contribute to Nociceptive Sensory Neuron Hyperexcitability Associated with Peripheral Neuropathies. J. Neurosci. 2019;39(8):1539–1550;DOI:10.1523/JNEUROSCI.0468-18.2018

27. Theile JW, Cummins TR. Inhibition of Navβ4 peptide-mediated resurgent sodium currents in Na_v_1.7 channels by carbamazepine, riluzole, and anandamide. Mol. Pharmacol. 2011;80(4):724–734;DOI:10.1124/mol.111.072751

28. Wengert ER, Saga AU, Panchal PS, et al. Prax330 reduces persistent and resurgent sodium channel currents and neuronal hyperexcitability of subiculum neurons in a mouse model of SCN8A epileptic encephalopathy. Neuropharmacology 2019;158:107699;DOI:10.1016/j.neuropharm.2019.107699

