## Supplemental Figures for "Functional and Pharmacological Evaluation of a Novel *SCN2A* Variant Linked to Early-onset Epilepsy"

### SUPPLEMENTAL INFORMATION

#### 1. Supplemental Figures

Fig. S1 Comparison between neonatal and adult M1879T channels

Fig. S2 Resurgent current response to carbamazepine

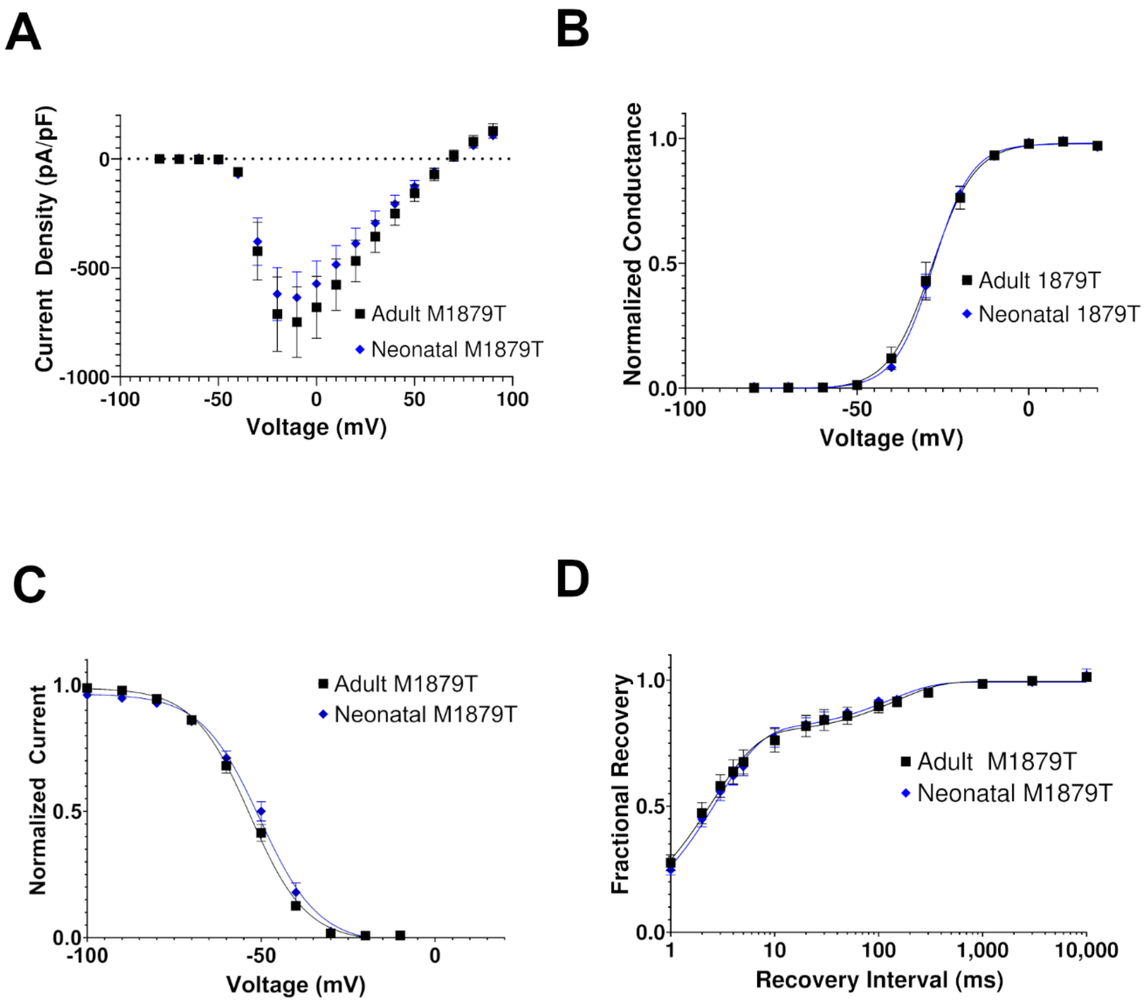


**Fig. S1. Comparison of functional properties between M1879T channels in either the neonatal of adult splice variant of SCN2A.**

(A) Current density-voltage plot of adult (n=8) and neonatal (n=15) isoforms of M1879T. (B) Voltage-dependence of activation for adult (n=14) and neonatal (n=12) isoforms of M1879T. No significant differences were observed between $V_{1/2}$ values (see Table 1). (C) Voltage-dependence of inactivation for adult (n=25) and neonatal (n=15) isoforms of M1879T. No significant differences were observed between $V_{1/2}$ values. (D) Logarithmic plot of the time course of recovery from inactivation comparing adult (n=10) and neonatal (n=11) isoforms of M1879T.

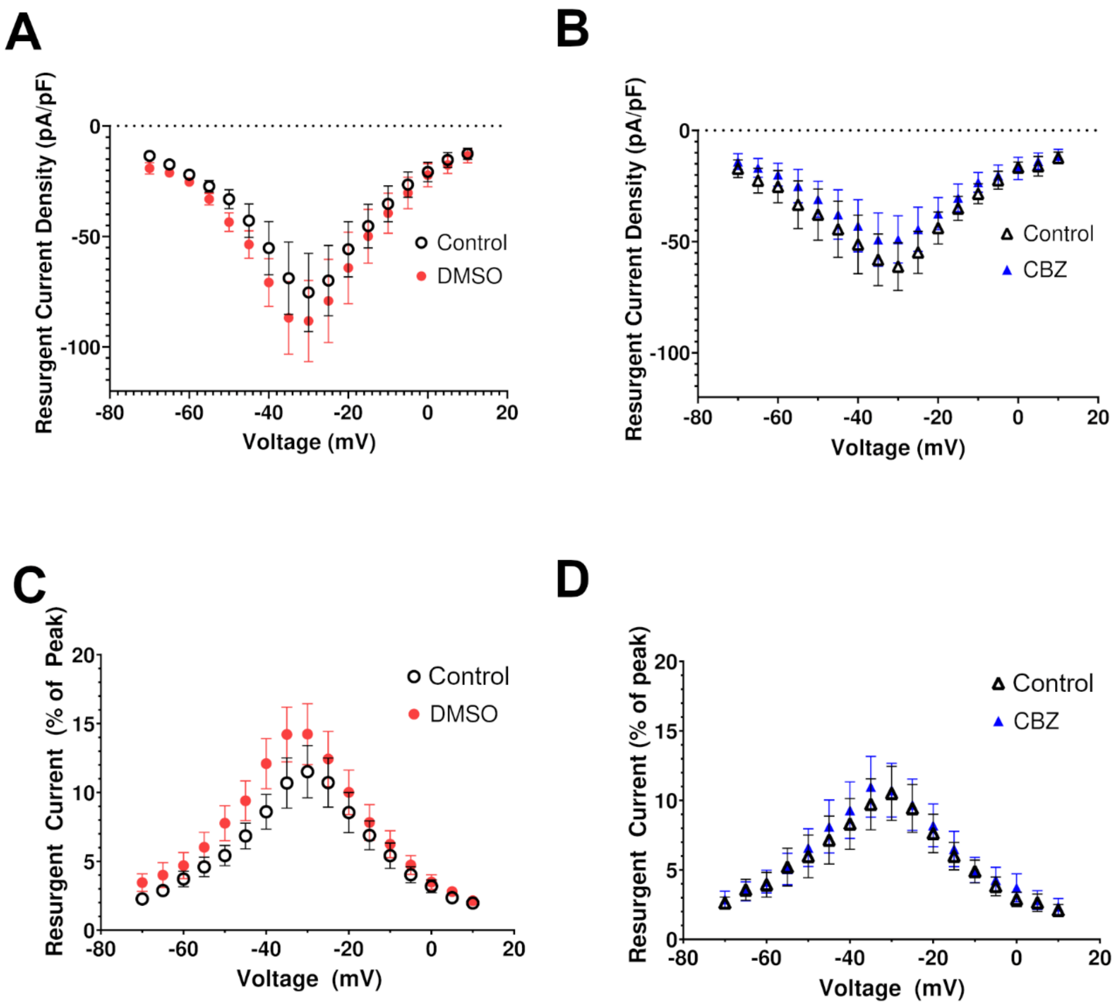


**Fig. S2. Resurgent current response to carbamazepine.**

(A) Voltage-dependence of resurgent current density in the control condition and after DMSO (vehicle) treatment (n=7). (B) Voltage-dependence of resurgent current density in the control condition and after 300 μM CBZ treatment (n=7). (C) Resurgent current as percentage of peak vs. voltage in control and after 300 μM CBZ treatment (n=7).
